# Using spatial prioritization to navigate current and future tradeoffs when controlling invaders

**DOI:** 10.64898/2026.05.29.728801

**Authors:** Brendan K. Hobart, H. Anu Kramer, Julianna M. A. Jenkins, Damon B. Lesmeister, Raymond J. Davis, M. Zachariah Peery

## Abstract

Controlling invasive species is a global conservation priority but is typically resource-limited, necessitating strategies to optimize control. Spatial prioritization methods can improve the efficiency of invader control by quantifying the benefits, costs, and risks of alternative intervention strategies. Yet prioritization designs are sensitive to the distinction between protecting native populations that are currently sympatric with invaders versus safeguarding presently allopatric native populations by preventing invader expansion. Gaining a better understanding of this distinction stands to improve our ability to prioritize the management of invaders whose distributions—and thus impacts on native species—are dynamic. We asked how prioritization designs addressing current versus future invader threats affected tradeoffs among focal species protection, biodiversity conservation, disturbance risk, and overlap with existing conservation infrastructure. As a case study, we spatially prioritized population control of invasive barred owls (*Strix varia*) in the northwestern US. We found that distinguishing between current versus future threats posed by barred owls to the native spotted owl (*S. occidentalis*) strongly mediated whether invader control stood to benefit native at-risk animal communities. Furthermore, this distinction also affected the degree to which population control would overlap with fire risk and federally protected forests, both of which plausibly affect the viability and success of conservation action. These results thus illustrate that deciding to prioritize the control of invaders based on their current versus future impacts on native species can dramatically affect the distribution and characteristics of high-priority areas for management. Our findings also directly inform control of barred owls in the northwestern US: we found that prioritizing future threats to spotted owls could protect at-risk amphibian communities from novel barred owl predation, but that high fire risk and minimal protected forest may complicate implementation. Thus, in both our system and more broadly, spatial prioritization methods are an important tool for quantitative, reproducible, and successful invader control.

## INTRODUCTION

From local preserves to international agreements, managing invasive species is a top conservation priority owing to the dramatic and irreversible ecological harm that invaders can precipitate (United Nations 2022, Roy et al. 2023). Yet, invader control (e.g., lethal removal) programs are often resource-limited and unable to implement comprehensive management, necessitating triage-like measures and calling for defensible strategies to select sites for control (Mack et al. 2000, Simberloff et al. 2005, Bottrill et al. 2008, Pyšek et al. 2020). Many schemes to optimize invader control incorporate information solely about the distribution and spread of invaders, implicitly assuming biodiversity and biogeographic conditions have uniform spatial distributions (Baker and Bode 2016, Nishimoto et al. 2021, Thompson et al. 2021). However, because species, habitats, and other factors are distributed unevenly across space, improved conservation outcomes can be achieved by leveraging spatially explicit biodiversity and biogeographic information when planning invader control (Januchowski-Hartley et al. 2018).

Spatial prioritization methods from the field of systematic conservation planning provide a quantitative and reproducible framework to optimize invader control efforts based on spatially variable patterns in biodiversity, costs, and risks (Januchowski-Hartley et al. 2011, Kukkala and Moilanen 2013, Giakoumi et al. 2025). Originally developed for designing protected areas, spatial prioritization methods have been applied in diverse ecological systems for a range of management goals, including invader control (Kukkala and Moilanen 2013, Januchowski-Hartley et al. 2018, Giakoumi et al. 2025). These applications demonstrate the value of spatial prioritization but highlight the need for a deeper understanding of synergies and tradeoffs when prioritizing invader control (Carter et al. 2024, Baker et al. 2025). Though many factors interactively affect the long-term success of invader control, several stand out as particularly important and prone to tradeoffs: invasion stage, threats to native species, disturbance risk, and existing conservation infrastructure. Better understanding how these factors alter optimal solutions for invader control and interact with each other can advance efforts to limit the ecological and economic harm of invasive species.

Invasion stage is a longstanding factor used by managers to help allocate limited resources to invader control efforts. It is well known that invasion stage mediates the effectiveness of invader control, with lower costs and greater chances of success early in an invasion, when invader population densities are low (and vice versa for late-stage invasions with high population densities) (Mack et al. 2000, Simberloff et al. 2013). Furthermore, when an invasive species’ range is expanding, the threat it poses to biodiversity is spatiotemporally dynamic—populations, species, and communities currently unaffected by invasion may become negatively impacted in the future following spread of the invader (Baker and Bode 2016). Critically, this spatiotemporal variability in threat means that optimal solutions for invader control may also shift over time. Although relatively little research has considered such possibilities, Carter et al. (2024) studied the current and future threat posed by an invasive willow, finding evidence for both win–wins and strong tradeoffs that depended upon whether current or future threats were prioritized. These findings point to the importance of quantifying tradeoffs when invader control is prioritized based on current versus future threats to native species (i.e., behind or beyond an invader’s leading edge, respectively), as doing so may reveal previously undiscovered synergies or risks.

Whether optimizing reserve design or invader control, tradeoffs in the protection of different taxa (i.e., “biodiversity coverage” in systematic conservation planning) are a common challenge (Kukkala and Moilanen 2013). In other words, management implementations that more comprehensively protect some species or groups often fail to protect others. Although systematic conservation planning aims to balance the protection of many species, conservation practitioners often struggle to generate data and interest for broad suites of species (Baker et al. 2019). For this reason, on-the-ground management—including invasive species control—is often motivated by and planned around the protection of key focal species (e.g., umbrella, flagship, or indicator species) (Simberloff 1998). A focal species approach can be simpler to operationalize but may also suffer more strongly from tradeoffs in biodiversity protection (e.g., if there are gaps in a focal species’ umbrella) (Carlisle et al. 2018). Within the context of using spatial prioritization to plan invasive species control, there is a need to better understand tradeoffs between the protection of focal species and broader biodiversity assets.

Invader control can also encounter tradeoffs involving disturbances, which can ultimately affect the implementation and success of management (Newton et al. 2011, Leroux and Rayfield 2014, Magris et al. 2015). Given that (a) many disturbance regimes are becoming increasingly novel (e.g., increasing in frequency and severity) (Turner and Seidl 2025), (b) novel disturbances threaten many native species (Devictor and Robert 2009, Legge et al. 2022, Ayars et al. 2023), and (c) invaders can often capitalize on recent disturbances (Hobbs and Huenneke 1992, Hierro et al. 2006, González-Moreno et al. 2015), explicitly testing for tradeoffs between optimal invader control and disturbance risk is likely essential. Identifying such tradeoffs can facilitate multiple paths towards improved invader control. For instance, spatial disturbance information (e.g., fire risk surfaces) can be included explicitly in prioritization exercises to identify areas that are optimal for invader control (e.g., areas of high biodiversity value) but have a low risk of being affected by disturbance (Hanson et al. 2025). Conversely, if all high-priority control areas also have high disturbance risk, invader control may need to be accompanied by preemptive measures that counteract disturbance-driven biodiversity impacts (e.g., fuels management to reduce severe fire) (Pressey et al. 2007).

Finally, invader control prioritizations may be improved by considering the spatial distribution of existing conservation infrastructure. Spatial prioritization methods can be leveraged to evaluate and encourage overlap between optimal management solutions and established conservation plans (e.g., protected areas), such that existing research, monitoring, and management infrastructure can contribute to new invader control efforts (Epanchin-Niell et al. 2010). For example, if areas deemed optimal for invader control occur within existing protected areas, invader control may gain support if it aligns with existing priorities and because consequent conservation gains may be relatively safeguarded against future invasions or other threats (Genovesi and Monaco 2013, Forner et al. 2022). Conversely, when locations outside of established conservation areas are targeted for invader control, managers may face threats of future development (Bertolino et al. 2021). Thus, improving our knowledge of tradeoffs involving biodiversity protection, disturbances, and existing conservation planning may help practitioners identify synergies and avoid risky or unworkable management plans.

Using spatial prioritization methods to inform invasive species control, we aimed to (1) evaluate tradeoffs among focal species protection, native species protection (biodiversity coverage), disturbance risk, and existing conservation infrastructure; (2) describe how such tradeoffs varied when current versus future threat was considered. As a case study, we prioritized population control of invasive barred owls (*Strix varia*) in the western United States (US), where this invader negatively affects an iconic focal species—the northern spotted owl (*Strix occidentalis caurina*)—and putatively impacts broader biological communities via novel competition and predation (Holm et al. 2016, Franklin et al. 2021, Dumbacher and Franklin 2024, Arenas-Viveros et al. 2025). In this system, we had four primary objectives: (1) Apply spatial prioritization to identify where barred owl population control would support federal policies to protect spotted owl populations. (2) Use systematic conservation planning approaches to quantify whether barred owl population control designed to protect spotted owls also affords benefits for other vertebrate species (i.e., herptiles, birds, and mammals) of conservation concern. (3) Examine whether tradeoffs exist between spotted owl protection, fire risk, and existing reserve infrastructure when prioritizing barred owl population control. (4) Determine whether prioritizing barred owl management based on current versus future threats to spotted owl populations modifies the outcomes of the above three objectives. Addressing these objectives can inform the control of a high-profile invasive species and advance our general understanding of how to prioritize such control efforts by providing example workflows and highlighting common pitfalls.

## METHODS

### Study system

We used spatial prioritization methods to study patterns in optimal population control of barred owls in the western US. Barred owls are native to the eastern US, but over the last century have spread to the western US via one or more anthropogenically-mediated pathways: tree planting and woody encroachment of the Great Plains and/or warming winters of southern Canada’s boreal forests (Livezey 2009a, 2009b). Barred owls have since established dense populations in western portions of Washington and Oregon, with the leading edge of their range now occurring in northern California (Jenkins et al. 2025). Via intense interference competition within forest habitats, barred owls now pose an existential threat to the federally threatened northern spotted owl throughout its range and likely pose a threat to the California spotted owl (*S. o. occidentalis*) in California’s Sierra Nevada (Franklin et al. 2021, Wiens et al. 2021, Hofstadter et al. 2022, Rockweit et al. 2023, Dumbacher and Franklin 2024). There is also cause for concern that barred owls are negatively impacting many other native species within these recipient ecosystems via novel competition and predation (Holm et al. 2016, Wiens et al. 2025). Indeed, relative to the spotted owls they are displacing, barred owls occur at higher densities and are far more generalized predators, consuming a wide breadth of vertebrates and invertebrates (Hamer et al. 2001, Kryshak et al. 2022, Baumbusch 2023, Arenas-Viveros et al. 2025). Owing to factors like habitat loss and fragmentation, many such prey were already species of conservation concern prior to the barred owl invasion, which may exacerbate population declines (Arenas-Viveros et al. 2025).

Driven by the conservation threat posed by barred owls, prioritizing and enacting control of their populations is a critical component of the northern spotted owl Recovery Plan (USFWS 2011), leading to research aimed at testing the efficacy of lethal removal to control barred owl populations (after deeming alternative methods like sterilization and relocation infeasible and ineffective) (Diller et al. 2014). Evidence has accumulated that lethal removals can diminish barred owl populations and stabilize spotted owl populations at landscape to regional scales (Diller et al. 2016, Wiens et al. 2021, Hofstadter et al. 2022), yet given the overwhelming size of the region, it is unlikely that comprehensive (i.e., wall-to-wall) removals can be achieved. Thus, this system offers a salient opportunity to use spatial prioritization methods to identify areas that are optimal for invader control and to test for the occurrence of tradeoffs among native species protection, disturbance risk, and existing conservation infrastructure.

We applied prioritization methods to the entirety of forested habitat within the northern spotted owl’s range in the US, which covers portions of California, Oregon, and Washington. Barred owl populations are dense and well-established in Oregon and Washington, whereas the invasion is at an earlier stage in parts of California, where population density decreases along a northwest-to-southeast gradient. Throughout this region, wildfire is currently the primary natural disturbance driving old forest loss (Reilly et al. 2021) and is a principal threat to spotted owl persistence alongside invasive barred owls (Jones et al. 2016, 2021). We thus focused on fire as a key disturbance which may inform or tradeoff with barred owl removals. The region also has a comprehensive conservation plan (the Northwest Forest Plan, NWFP; (USDA and USDI 1994)) that, among many actions, sets aside forested land within the range of the northern spotted owl for conservation, maintenance, and restoration of late-successional/old-growth forest. These areas, broadly designated as “federally reserved forest” in the NWFP, offer areas where barred owl control may be the most beneficial for native species associated with older forests.

### Data compilation

We compiled a suite of spatial data to capture barred and spotted owl landscape use, threatened species diversity, fire risk, federally reserved forest, and other background factors (Fig. 1A). Regardless of their native resolution and alignment, all spatial data were resampled and aligned to a “master raster” with 30 m resolution. The master raster extent was defined as the area with complete coverage for all input spatial datasets across forested land within the NWFP boundary.

**Figure 1.**
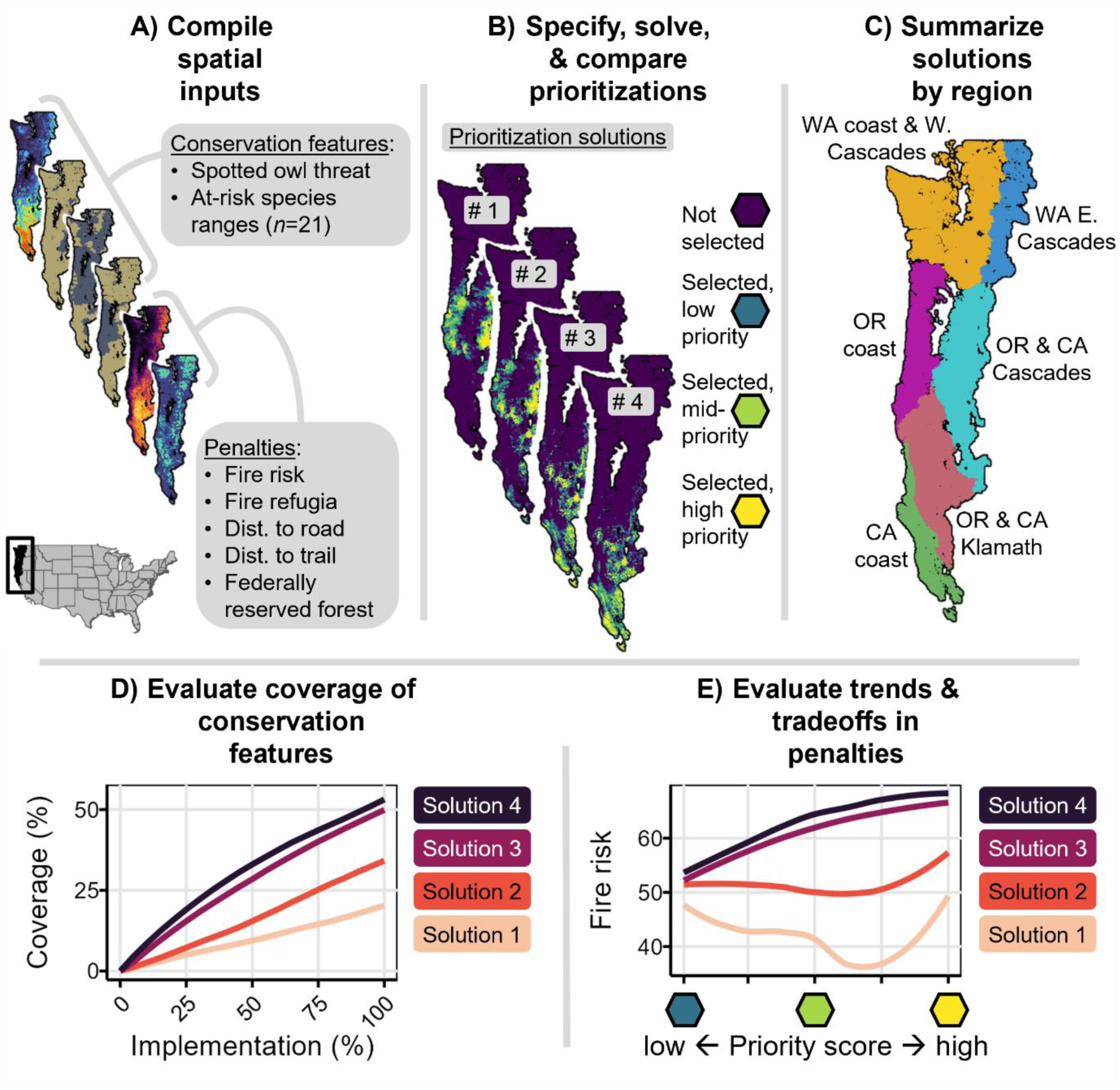
Conceptual workflow of our research on prioritizing the spatial allocation of barred owl (*Strix varia*) population control in the northwestern United States. (A) We compiled, aligned, and summarized input spatial data within 5 km^2^ hexagonal cells, before (B) being included in prioritization analyses that were solved (i.e., optimized) under a range of weight specifications. After solving and mapping prioritizations, we (C) summarized the distribution of high-priority hexagonal cells among regions in the study area, (D) quantified coverage of at-risk species by prioritizations, and (E) evaluated the distribution of penalties among solutions.

To capture barred and spotted owl distributions across the region, we compiled data from Jenkins et al. (2025) and Lesmeister et al. (in review), who used occupancy models to analyze acoustic monitoring detection data collected in 2023. Barred owl landscape use and spotted owl pair occupancy (both are hereafter referred to as “landscape use” for brevity) were modelled as functions of various ecological and biogeographical variables. Occupancy model results were then used to predict contemporary region-wide barred and spotted owl landscape use (see Jenkins et al. (2025) for full methodological details). Importantly, because their models were fit to data collected in 2023—after the abundance and distribution of spotted owls was altered by invasive barred owls across much of the former’s range—predicted spotted owl landscape use reflects anticipated landscape use after accounting for competitive exclusion by barred owls.

To quantify the spatial distribution of at-risk biodiversity (i.e., non-spotted owl species), we adopted a list originally compiled for the USFWS Barred Owl Management Strategy Environmental Impact Statement (Table 3-29 in US Fish and Wildlife Service [2024a]). The original list contained those species of conservation concern (federally listed, state listed, and/or special concern status) that are putatively impacted by invasive barred owls via competition or predation. We filtered the original list to include only terrestrial species that had geographic ranges and geographic range data within our study area. The final list included 21 species: 4 birds (e.g., the marbled murrelet [*Brachyramphus marmoratus*]), 9 mammals (e.g., the Western grey squirrel [*Sciurus griseus*]), and 8 herptiles (e.g., the foothill yellow-legged frog [*Rana boylii*]; see Table S1 for the full list of species and their listing statuses). For each species on our list, we obtained a shapefile of their geographic range from the US Geological Survey (USGS) Gap Analysis Project (GAP; https://www.usgs.gov/programs/gap-analysis-project), which was transformed into a binary raster aligned with the master raster described above.

We estimated contemporary fire risk using the output of analyses from Davis et al. (2017). Briefly, Davis et al. (2017) mapped the suitability of large forest fires (>40 ha) within our study area by modelling recent fire occurrence (1971–2000) as a function of precipitation, temperature, slope, and elevation, making spatial predictions of fire suitability, and testing these predictions against an independent fire occurrence dataset (from an independent time period: 2001–2015). For full methodological details, see Davis et al. (Davis et al. 2017).

We used the NWFP Land Use Allocation (LUA) map to summarize the extent of federally reserved forest across the region. Federally reserved forest (under the NWFP) is a broad classification that occurs on lands administered by any federal agency (i.e., US Forest Service, National Park Service, and Bureau of Land Management) and includes categories such as congressionally reserved areas (i.e., wilderness, national parks and monuments, etc.), late-successional/old-growth forest reserves, riparian reserves, and others (Davis et al. 2011, 2022). Although reserves designated by other entities (e.g., states or NGOs) occur throughout the region, federally reserved forest is highly relevant to barred owl management, given the recent development of the Barred Owl Management Strategy by the US Fish and Wildlife Service, who will also provide oversight of barred owl management.

Finally, we gathered spatial data for several factors that were included in prioritizations but were not the focus of our analyses. These included fire refugia, distance to roads, and distance to trails. Fire refugia are relevant to management planning in this region because they regularly burn at lower severities than surrounding landscapes and likely represent historical (i.e., pre-fire suppression) spotted owl nesting and roosting habitat (Lesmeister et al. 2025). We estimated fire refugia with maps from the Eco-Vis portal (Naficy et al. 2021), which uses established methods (Meigs et al. 2020, Downing et al. 2021) to predict the occurrence of fire refugia throughout the northwestern US. We used the contemporary (2020) topoclimatic fire refugia product from Eco-Vis, which is based on the results of statistical models that relate the occurrence of fire refugia to topography and climate.

Trails and roads provide means for managers to access mountainous forests in our study region and are thus key drivers of logistical feasibility. We calculated distance to road and distance to trail by processing the National Transportation Dataset of The National Map (U.S. Geological Survey 2023) using the ‘Euclidean Distance’ tool within the Spatial Analyst toolbox of ArcGIS (ESRI; Redlands, CA). To minimize potential edge effects, we calculated the distance to the nearest road or trail within the 50 km buffered study area and then clipped this layer to the study area.

### Defining current and future threat to spotted owls

Building on the methods of Carter et al. (2024), we combined spatial data on barred and spotted owl landscape use to formulate two layers: one that captured the current threat posed to spotted owl populations by barred owls, and one that captured the future threat posed to spotted owl populations by barred owls. Conceptually, the current threat posed to spotted owls by barred owls is greatest where both species co-occur at relatively high densities; in other words, where spotted owls persist behind the barred owl invasion front. On the other hand, the greatest future threat occurs near or beyond the invasion front, in areas where spotted owls have experienced less competitive exclusion, but barred owls will eventually invade (or increase in density) if left unchecked. Thus, on a per-pixel basis, we calculated current threat as the product of spotted and barred owl landscape use probabilities:

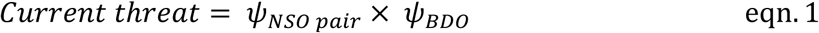

where *ψNSO pair* denotes the probability of landscape use (after accounting for imperfect detection and environmental covariates) of a pair of northern spotted owls and *ψBDO* denotes landscape use probability for any barred owl (Jenkins et al. 2025). Conversely, to calculate future threat, we subtracted barred owl landscape use probability from spotted owl landscape use probability:

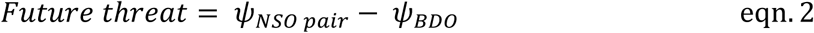

These formulations resulted in high current threat values for pixels where both barred and spotted owl landscape use probabilities were high, and high future threat values in pixels where spotted owl landscape use probabilities were high but barred owl landscape use probabilities were low (Fig. 2). Formulating future threat in this way made the reasonable assumption that in the absence of management, barred owls will disperse to virtually every portion of the northern spotted owl’s range. In California, we have observed barred owls as far southwest as Marin County and as far southeast as the central Sierra Nevada (Wood et al. 2021, Hofstadter et al. 2022, Hobart et al. 2025). Thus, relatively healthy spotted owl populations currently occurring in areas with few or no barred owls will presumably face invasion and increased interspecific competition in the future if no population control of barred owls is carried out.

**Figure 2.**
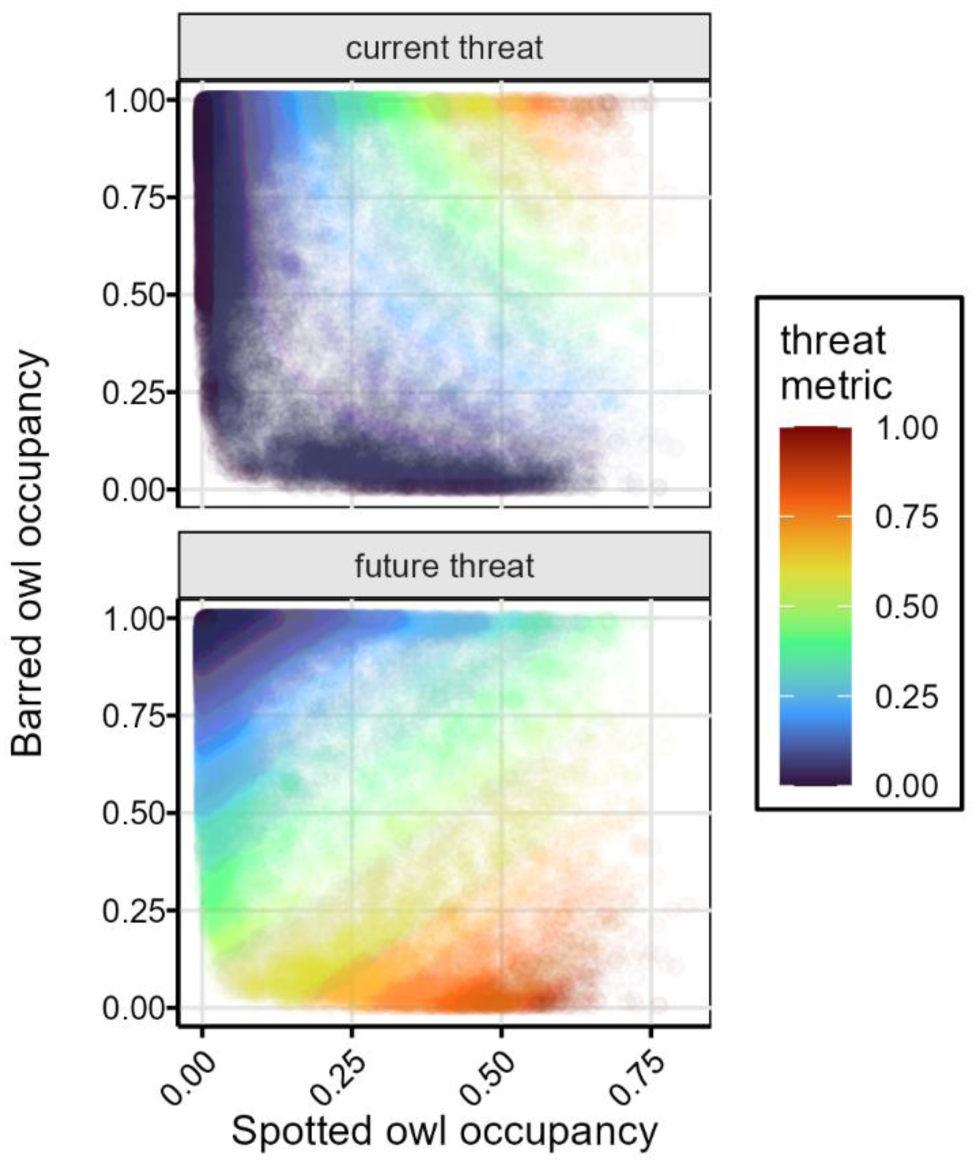
To better capture and quantify spatiotemporal variability in threat posed to northern spotted owls (*Strix occidentalis caurina*) by invasive barred owls (*S. varia*) in the northwestern United States, we developed two metrics of threat. Current threat (top) had maximum values in areas with high predicted landscape use of both spotted and barred owls; this was motivated by the fact that extant spotted owl populations overlapping with barred owls face an immediate risk of extirpation from competition. Conversely, future threat (bottom) had maximum values in areas with high predicted landscape use of spotted owls but low predicted landscape use of barred owls, which will expand eventually to have complete range overlap with northern spotted owls.

### Spatial prioritization

To designate planning units across which prioritizations would be conducted, we selected a grid of 5 km^2^ hexagonal cells that is widely used across the NWFP region for management and monitoring (Lesmeister et al. 2021). To establish inclusion of cells in prioritization analyses, we summarized coverage of the master raster in each hexagonal cell; cells with <25% coverage by the master raster were omitted, yielding a study area consisting of 148,192 hexagonal cells (740,960 km^2^). Next, within each hexagonal cell, we summarized all spatial data inputs by taking their mean value across all 30 m pixels whose centroid was inside the hexagon. Finally, to improve the selection of weight parameters (for conservation features and penalties), we rescaled each hex-level variable to have a minimum of 0 and a maximum of 1.

We used the ‘prioritizr’ package (Hanson et al. 2025) in R (R Core Team 2019) to define prioritization problems and solve them to optimize the spatial allocation of barred owl removal efforts (Fig. 1B). The prioritizr package provides a platform for optimizing the spatial allocation of conservation actions such as protected area designation, habitat restoration, and reintroductions. Given the relevance of spotted owls to conservation planning in the region, our primary focus was varying the weight assigned to spotted owl threats, which we denote *W_NSO_* (for “northern spotted owl weight”). In two separate series of prioritizations, we varied the weight assigned to layers capturing current and future threat to spotted owls, respectively. Preliminary sensitivity analyses revealed that varying *W_NSO_* between 1–15 captured approximately the minimum and maximum influence of spotted owl threat on optimal solutions. Thus, we fixed *W_NSO_* at values of 1, 5, 10, and 15. Weights for all other spatial inputs were held constant at positive or negative one to prioritize for or against these layers: geographic ranges for all 21 at-risk species (Table S1), fire risk, federally reserved forest, fire refugia, distance to road, and distance to trail. Species ranges were included as conservation features with weights fixed at positive one. All other layers were included as penalties with weights fixed at positive one (i.e., penalizing: fire risk, distance to road, and distance to trail) or negative one (i.e., incentivizing: federal reserved forest and fire refugia). See Table S2 for weighting values summarized across all prioritization iterations.

When specifying prioritization problems, we used the “maximum utility” objective function, which maximizes coverage of input layers (while accounting for layer-specific weights) given some user-specified budget. We defined cost per planning unit as its land area (i.e., 5 km^2^) and specified a budget of 45,000 km^2^ (i.e., 9000 hexagonal cells) per solution, which represents roughly the maximum land area across which barred owls can be managed under the USFWS Barred Owl Management Strategy (US Fish and Wildlife Service 2024b). Once fully specified, we used prioritizr to optimize the allocation of barred owl removal effort across hexagonal cells within the study area based on the distribution and weighting of input layers. We used the commercial Gurobi exact algorithm solver (Gurobi Optimization LLC 2020) to implement optimizations, although open-source solvers can also be used (Hanson et al. 2025). Finally, following optimization, we called prioritizr’s “eval_rank_importance” function, which assigns a relative importance score (ranging from zero to one) to each unit (hexagonal cell) that is selected for management. Briefly, the function implements methods developed by Jung et al. (Jung et al. 2021), in which the prioritization problem is re-solved across a range of evenly spaced, decreasing budgets (we specified 20 iterations). Only those planning units (i.e., hexagonal cells) selected for management in the first optimization are considered viable for inclusion in the second optimization, and so on, such that each subsequent solution contains a subset of previously selected units. Finally, the proportion of iterations in which a given unit is selected is taken to represent that unit’s relative importance, or, in other terms, its cost-effectiveness for meeting the selected objective.

After optimizing each prioritization scenario and calculating importance scores, we retrieved the output table in which each hexagonal cell was designated as being selected for management and, if it was, its relative importance score. We first joined this table with geographic data for each hexagonal cell, and mapped solutions across different scenarios (i.e., current versus future threat) and *W_NSO_*. We summarized the distribution of high-priority cells (i.e., those with relative importance values >0.9) across biogeographic regions of our study area (Fig. 1C), defined using the NWFP modelling regions (Davis et al. 2022).

Next, we joined the optimized solutions with the input spatial data to evaluate coverage (Fig. 1D), which captures the extent of a given layer’s distribution that is covered by a given solution (Hanson et al. 2025). To quantify coverage, we reported the amount of each species’ distribution that was captured by each solution across the gradient of relative importance scores. For all hexagonal cells assigned a given relative importance score (distributed between zero and one in 20 evenly spaced increments), we summed each feature’s coverage, then calculated the cumulative sum across relative importance values, moving from high to low (under the assumption that if incomplete solution implementation occurs, it would first incorporate cells with higher relative importance). For species other than the spotted owl, we averaged these cumulative coverage curves across all species within a given taxonomic group: birds, mammals, and herptiles (species-specific coverage curves are provided in Fig. S1).

Finally, to evaluate trends and tradeoffs in fire risk and federally reserved forest, we took the mean value of each layer across all hexagonal cells receiving a given relative importance score for a given solution (Fig. 1E). We then used generalized additive models (implemented with the ‘geom_smooth’ function from the ggplot package (Wickham 2011)) to fit a smoothed curve to the distribution of mean values across relative importance scores.

## RESULTS

Across our study area, the geographic location of cells deemed high priority (relative importance score >0.9) for barred owl management varied with *W_NSO_* (the weight assigned to spotted owl threat when specifying conservation problems) and whether current or future threat was prioritized (Fig. 3). Under both current and future threat scenarios, low *W_NSO_* (i.e., weight equally allocated to NSO threat and all other at-risk species) resulted in most high-priority cells occurring in the Oregon and California Cascades region (74–78%), with fewer cells in the Oregon Coast Range and Klamath regions (13–17% and 9%, respectively; Table S3). In the scenario where current threat was considered and *W_NSO_* was increased to its maximum value (*W_NSO_*=15), the distribution of high-priority cells shifted primarily to the Klamath region (79%), with some cells remaining in the Oregon and California Cascades region (15%), and few cells shifting into the California Coast Range region (6%; Table S3). Conversely, when *W_NSO_* was increased under the future threat prioritization, the distribution of high-priority cells shifted further south and west: at maximum *W_NSO_*, high-priority cells were roughly evenly divided between the Klamath and California Coast Range regions (52% and 47%, respectively), with <1% of high-priority cells occurring in the Oregon and California Cascades region. Notably, aside from the Olympic Peninsula—which contained a modest proportion (3%) of moderate-priority cells (relative importance score 0.5–0.9) when prioritizing current threat (*W_NSO_*=15)—regions in the state of Washington contained no moderate-to-high priority cells under any scenario.

**Figure 3.**
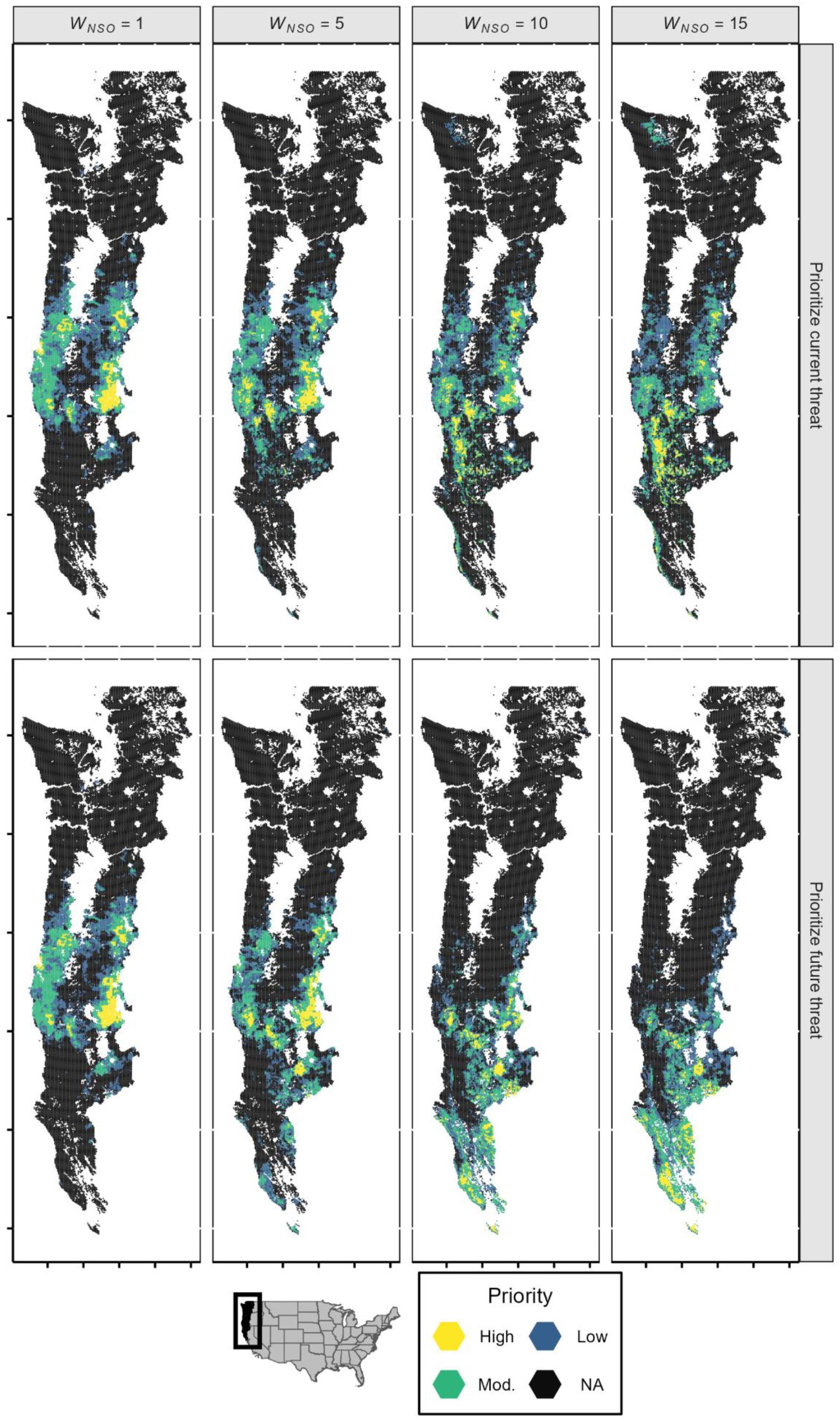
Maps depicting the location and importance of areas selected for invasive barred owl (*Strix varia*) population control during prioritization analyses. Each 5 km^2^ hexagonal pixel across the study area has been colored according to its status for each prioritization. Cells shaded dark gray denote those not selected for barred owl control during prioritization. Conversely, colored cells were selected, with increasingly bright colors denoting increasing relative importance. Relative importance scores (which vary continuously between 0 and 1) were binned into three categories: 0–0.5 (low), 0.5–0.9 (moderate), and 0.9–1.0 (high). The weight assigned to threats posed to northern spotted owls (*S. occidentalis caurina*) by barred owls (*W_NSO_*) varies among columns, whereas the temporal specification of threat varies among rows (i.e., current versus future threat).

Across spotted owls and 21 other at-risk native species (Table S1), the degree of protection (i.e., coverage) afforded by prioritization solutions varied with *W_NSO_*, whether current or future threat was prioritized, and the taxa being considered (Fig. 4). As expected, cells with relatively high spotted owl landscape use were covered more completely when scenarios had a higher specified *W_NSO_* value. Notably, when prioritizing future threat, increasing *W_NSO_* led to increases in the coverage of at-risk amphibians: if fully implemented, the maximum-*W_NSO_* scenario afforded on average 26% greater coverage of amphibian ranges relative to the lowest-*W_NSO_* scenario (Fig. 4). Conversely, when current threat was considered, increasing *W_NSO_* had little effect on the coverage of at-risk amphibians. Across scenarios considering both current and future threats to spotted owls, coverage of at-risk birds declined as *W_NSO_* increased. At full implementation of scenarios considering current and future threat, the coverage of at-risk bird species declined by 10% and 17% respectively. Finally, there were only modest changes in the coverage of at-risk mammals across different prioritization scenarios.

**Figure 4.**
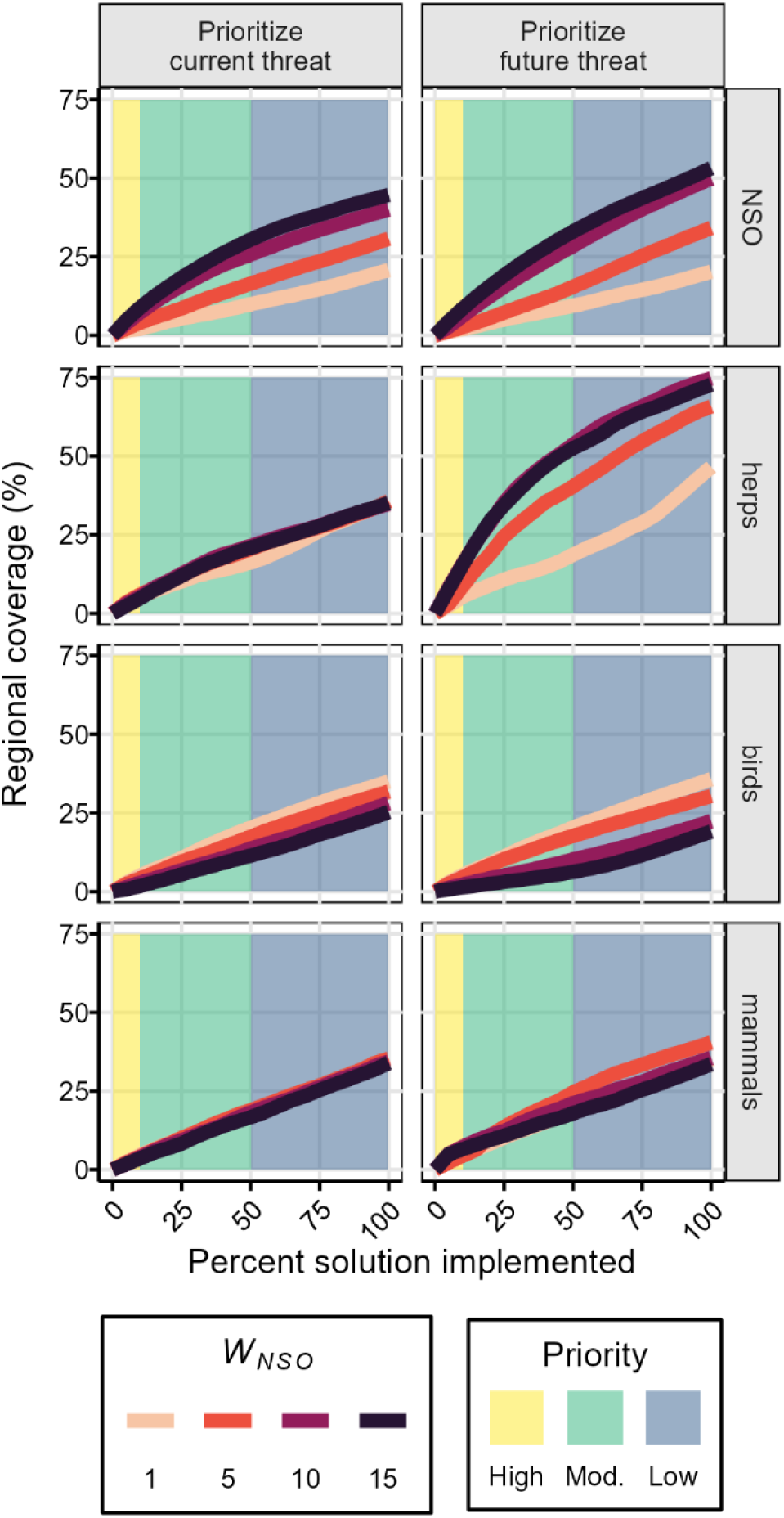
Results from coverage analysis of our prioritizations for population control of invasive barred owls (*Strix varia*) in the northwestern US. Prioritizations were largely organized around addressing the threat posed to native northern spotted owls (*S. occidentalis caurina*) by barred owls. We varied the importance (weight) assigned to this threat (denoted *W_NSO_* and indicated by line color), and fit prioritizations based on both current and future threats (left and right columns, respectively). Curves show the coverage of each prioritized solution for different native, at-risk taxonomic groups (each row). A curve that rises more steeply and to a greater maximum is indicative of a more efficient solution—greater coverage is afforded with relatively less effort (that is, less of the solution being implemented). Background shading denotes priority bins and is aligned with the shading in Figure 3.

The distributions of disturbance risk (fire suitability) and conservation infrastructure (federally reserved forest) varied widely across prioritization scenarios (Fig. 5). Whether prioritizing current or future threat, increased *W_NSO_* values were associated with greater fire suitability values. This was especially true when prioritizing future threat: fire suitability was high (67–73 on a unitless scale from 0–100) across moderate and high priority cells (i.e., relative importance score > 0.5) when *W_NSO_* was high (i.e., held at 10 and 15). On the other hand, when prioritizing current threat, fire suitability values rose only for the highest priority cells (i.e., areas in the Klamath region with high current threat also had high fire risk). When current threat was prioritized, high priority cells (relative importance score >0.9) had high percentages of federally reserved forest (56–84%), regardless of *W_NSO_*. Conversely, when future threat was prioritized, increasing *W_NSO_* values were associated with declining federally reserved forest: within high priority cells, the average hex-level percentage of land designated as federally reserved forest declined precipitously from 61–71% at low *W_NSO_* values (*W_NSO_* = 1, 5) to 35–38% at high *W_NSO_* values (*W_NSO_* = 10, 15).

**Figure 5.**
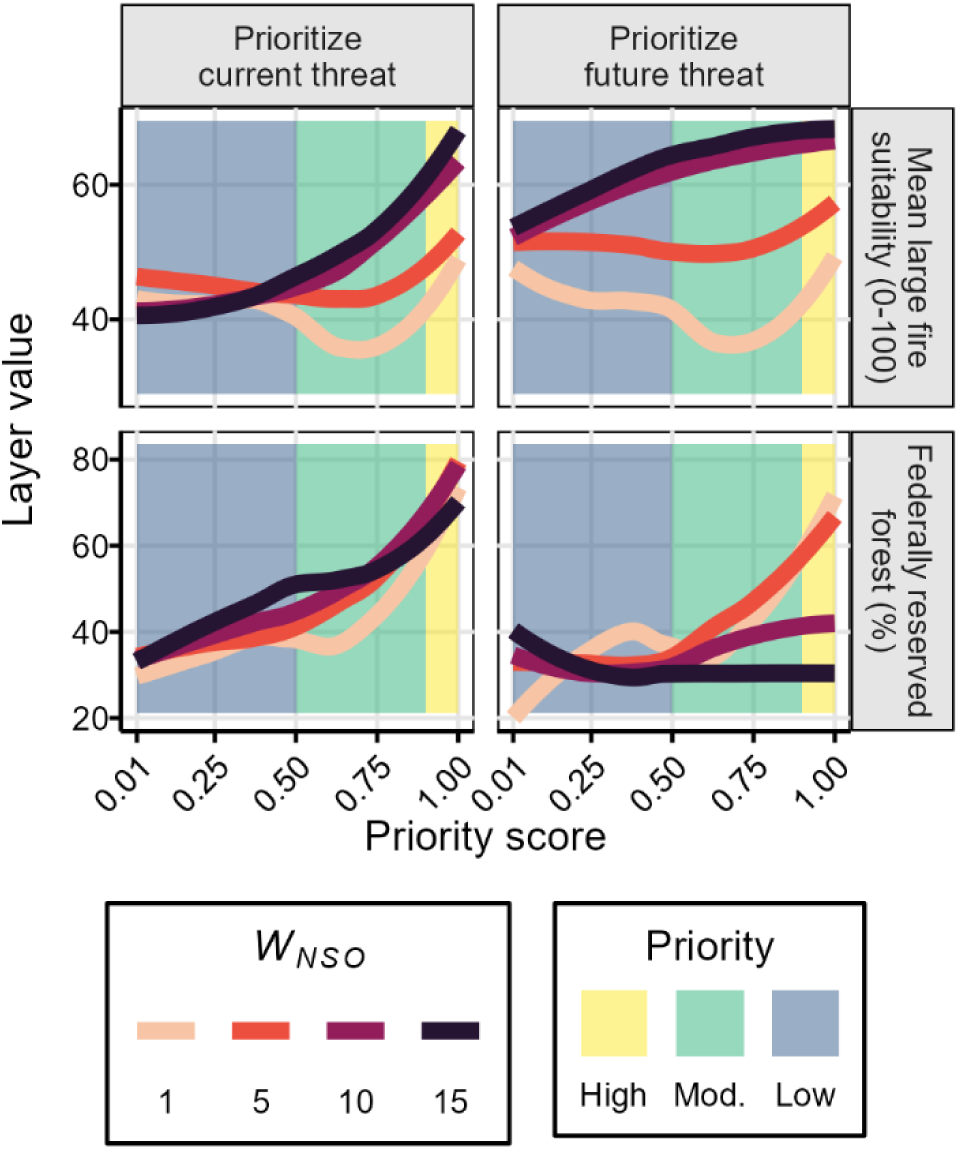
(penalties) Smoothed distributions of fire risk (top row) and federally reserved forest (bottom row) across areas of low-to-high priority for controlling barred owl (*Strix varia*) populations. We fit prioritizations to address both current (left column) and future (right column) threats posed to native northern spotted owls (*S. occidentalis caurina*) by barred owls, with the weight assigned to these threats (denoted *W_NSO_*) indicated by line shading. We calculated mean layer values across all hexes within equal-interval priority score bins, then used basic general additive models to plot smoothed lines to mean values across priority scores. Background shading denotes priority bins and is aligned with the shading in Figure 3.

## DISCUSSION

Implementing management to control invasive species is an effective and pressing conservation action (Langhammer et al. 2024), yet it is almost always resource limited (Simberloff et al. 2005). Here, adopting methods for spatially optimizing invader control, we advanced knowledge of tradeoffs that arise during such implementation, which stands to improve the efficiency and efficacy of invader control efforts. As a case study, we used prioritization methods to understand the intricacies of planning invasive barred owl population control in the northwestern US. We found that organizing prioritizations around the protection of a focal species, the northern spotted owl, modified the spatial distribution of and tradeoffs associated with high-priority areas for barred owl control, and that prioritizing current versus future threats further modified outcomes. Collectively, our work highlights that when prioritizing and planning invasive species control, incorporating information on the spatiotemporal variability in threats posed to native species can reveal both novel opportunities and unforeseen complications.

Planning invader control around the protection of focal (e.g., umbrella or indicator) species may allow conservation practitioners to leverage policy and funding opportunities but requires careful consideration (Simberloff 1998). Given the relevance of spotted owls for management and policy in the northwestern US (Spies et al. 2019), we organized and specified our prioritizations around addressing the threat posed to this focal species by invasive barred owls. Increasing the weight assigned to spotted owls (*W_NSO_*) notably shifted prioritization outcomes, such that there was limited overlap in the location of high-priority cells between spotted owl-focused prioritizations (i.e., *W_NSO_* = 15) and baseline prioritizations (i.e., *W_NSO_* = 1), where the latter represented a scenario aimed at protecting a broad suite of at-risk species. Furthermore, at maximum weight assigned to spotted owls (i.e., *W_NSO_* = 15)—which is representative of federal priorities in the region (US Fish and Wildlife Service 2024b)—large shifts in the geography of high-priority areas for barred owl control were driven by the decision to prioritize current versus future threats. This finding reinforces that it is important to consider temporal variation in invader impacts when making decisions about control efforts (Carter et al. 2024).

For invaders with expanding ranges (including barred owls), there are meaningful consequences of choosing whether to address current or future threats to native species. Indeed, extensive research on invasion stage shows that these alternatives require managers to adjust intervention strategies and locations (Simberloff et al. 2013). For example, in Washington, where the barred owl invasion is longstanding and spotted owl densities are low (Lesmeister et al. in review, Jenkins et al. 2025), our results indicate that barred owl population control is not high priority (relative to elsewhere in the region), suggesting that alternative conservation actions (e.g., captive breeding of spotted owls) may be more appropriate (Dumbacher and Franklin 2024). Further south, in the Klamath region of Oregon and California, prioritizing control of barred owl populations based on our current threat metric may prevent extirpations of spotted owls, albeit while entailing costly, intensive efforts owing to the high density of established barred owl territories in this region (Jenkins et al. 2025). On the other hand, choosing to prioritize control using our future threat metric may fail to prevent such extirpations, but would safeguard relatively healthy spotted owl populations in the southern Klamath region and California Coast Range with relatively less effort because action could be taken earlier in the invasion process (Hofstadter et al. 2022, Watson et al. 2023, Hobart et al. 2025). In the case of barred owls and in general, mitigating both current and future threats posed by invaders would be ideal, but resource limitations preclude doing so in many systems. Because invasion stage is key, allocating resources to control expanding invader populations proactively or reactively is fundamental to invasion biology yet is only rarely considered in spatial prioritization (Carter et al. 2024). Our work shows that explicitly incorporating the temporal dimensions of invader impacts into prioritizations can reveal both opportunities for efficient biodiversity conservation and unforeseen tradeoffs.

We found that prioritizing invader control based on threats posed to a focal species could confer protection to other at-risk species, but that such benefits were variable among taxa and shifted when current versus future threats were prioritized. Coverage analysis of our spatial prioritizations revealed that relative to baseline scenarios (*W_NSO_* = 1), those with high weight assigned to future spotted owl threat provided considerably greater average geographic coverage for at-risk herptile species—most of which were amphibians (*n* = 7). This finding aligns well with previous research in northern California that similarly suggested spotted owl-based conservation benefits native salamander species (Dunk et al. 2006). Given that amphibians are declining in the northwestern US (Luedtke et al. 2023) and are extensively consumed by barred owls (Kryshak et al. 2022, Arenas-Viveros et al. 2025), such findings point to an important possible synergy whereby controlling barred owl populations for the benefit of spotted owls could help protect diverse and at-risk amphibian communities.

Our application of a focal species approach for planning invader control also revealed conservation pitfalls, notably that average coverage of at-risk bird species declined when future spotted owl threat was highly weighted. Although this effect was comparatively modest, it is worth noting that relatively few bird species (*n* = 4) were deemed at-risk under the current framework—the addition of new species based on recent research (e.g., Rugg et al. 2023, Wiens et al. 2025) and updated conservation plans (e.g., State Wildlife Action Plans) may shift such outcomes. Regardless, this outcome highlights the reality that in our system, and more broadly, it may not be plausible to protect all taxa equally from invader impacts. Such tradeoffs are common in conservation planning and are likely exacerbated by approaches that hinge upon focal species with somewhat specialized umbrellas (Brunk et al. 2025), as in the case of spotted owls and mature forest species in the northwestern US. Despite this challenge, we show how applying prioritization methods to the planning of invader control can make explicit both the tradeoffs at hand and how they are affected by management decisions (e.g., addressing current or future threat).

Although prioritization applications tend to focus on quantifying and understanding biodiversity protection, these methods also lend valuable insights into tradeoffs with abiotic and logistical factors that arise when planning invader control programs. We found that changing prioritization specifications had a large impact on the degree to which high-priority cells overlapped with disturbance risk (fire suitability) and existing conservation infrastructure (federally reserved forest). These results add important nuance to the interpretation of the biodiversity-focused outcomes discussed above. Notably, although prioritizing future threats to spotted owls could yield relatively efficient control of barred owls and simultaneous benefits to herptiles, such solutions were also associated with high fire risk and low federally reserved forest. Thus, if managers wish to capitalize on the benefits of prioritizing future threats, they will have to contend with possibly catastrophic wildfires and complex sociopolitical landscapes. This suggests there may be value in pairing invader monitoring and control with forest restoration aimed at reducing the probability and severity of future wildfires (e.g., fuels treatments). Although such restoration would incur additional costs, our results highlight that reducing the risk of wildfires may be a prerequisite to realizing the benefits of controlling barred owls—clearing landscapes of this invader will contribute little to the conservation of forest species if their habitat is subsequently destroyed by fire.

Beyond heightened fire risk, limited overlap of high-priority cells (under the future threat prioritization) with federally reserved forest points to additional challenges, but also potentially opportunities. Under most future threat scenarios, this pattern was driven by high prioritization scores in the California Coast Range region, where large areas of private timberlands have little federally reserved forest but varying degrees of suitable spotted owl habitat (Diller et al. 2012). Limited overlap between high-priority cells and federally reserved forest suggests that implementing barred owl population control based on future threats to spotted owls will require cooperation among landowners and management agencies (Epanchin-Niell et al. 2010). This presents a perceived hurdle to barred owl control, given that effective conservation (whether invader control or forest restoration) in landscapes with complex ownership is generally challenging, requiring buy-in and commitment from diverse stakeholders (Burger et al. 2019). Yet, in our study system, buy-in from private stakeholders has already been partially achieved: land-owning timber resource companies have been at the forefront of studying how to control barred owl populations in northwestern California (Diller et al. 2014, 2016) and Habitat Conservation Plans (HCPs) have been leveraged to codify and incentivize extensive barred owl control by private entities (Green Diamond Resource Company and US Fish and Wildlife Service 2019, Sierra Pacific Industries and US Fish and Wildlife Service 2020). If population control can be balanced with the retention of critical habitats and land use activities, similar incentive structures might more broadly present opportunities for conducting invader control in high-priority areas, despite complex ownership patterns.

Furthermore, the implementation of potential forest restoration practices (discussed above) may be difficult to achieve in landscapes with extensive formal restrictions on timber harvest in mature forests (but see Lesmeister et al. 2025). For example, in the decades since the NWFP’s development, fire-resiliency restoration activities have been lacking within areas designated as federally reserved forest (Spies et al. 2019). Thus, while our prioritization design reflected the notion that high overlap with federally reserved forest is generally a desirable feature of a federally overseen barred owl control plan (US Fish and Wildlife Service 2024b), lands outside federally reserved forest may offer unique opportunities to implement and sustain barred owl population control. Although the strength and context of tradeoffs involving invader removal, disturbance risk, and existing conservation infrastructure certainly vary among systems, such tradeoffs are likely common and, as shown here, can be formally explored and quantified using spatial prioritization.

We have shown how prioritization methods can be used to optimize invader control, understand tradeoffs among relevant factors, and explore the influence of temporal variation in threats to biodiversity. Our results also hold direct relevance to implementing population control of invasive barred owls in the northwestern US, where this species threatens myriad native species (Holm et al. 2016, Arenas-Viveros et al. 2025). Despite strong evidence that controlling barred owl populations can successfully benefit native species (Wiens et al. 2021, Hobart et al. 2025), such control has spurred considerable controversy (Dumbacher and Franklin 2024), driving the need to identify approaches that simultaneously maximize biodiversity protection while minimizing the number of barred owls killed. To this end, our analyses offer a valuable and novel distinction—prioritizing removals based on current versus future threats—that clarifies the benefits and complexities of managing barred owl populations in a proactive versus reactive manner. Like all spatial prioritization outcomes, our results should be viewed as information capable of supporting management decisions, rather than data upon which decisions can be made outright. For instance, spatial prioritization outcomes may be combined with expert opinion or up-to-date occurrence records of protected native species to help identify areas for invader control (Hanson et al. 2025). Thus, when applied alongside existing methods for planning invasive species control, spatial prioritization stands to improve efficiency, reproducibility, and accountability, all of which are essential to addressing the threat that invasive species pose to biodiversity.

## Acknowledgements

We are thankful to the many scientists and practitioners who have developed protocols and plans for addressing invasive barred owls in the northwestern US. For their contributions in the form of engaging conversations, we especially thank D Hofstadter, R Bown, and K Fitzgerald. We are grateful to the NASA Biological Diversity & Ecological Conservation program for funding this work. The findings and conclusions in this publication are those of the authors and should not be construed to represent any official U.S. Government determination or policy. The use of trade or firm names is for reader information only and does not imply endorsement by the U.S. Government.

## Author Contributions

BKH: conceptualization, methodology, software, formal analysis, investigation, data curation, writing – original draft, writing – review & editing, visualization, project administration.

HAK: methodology, software, data curation, writing – review & editing.

JMAJ: methodology, data curation, writing – review & editing.

DBL: methodology, data curation, writing – review & editing.

RJD: methodology, data curation, writing – review & editing.

MZP: conceptualization, investigation, resources, writing – review & editing, supervision, project administration, funding acquisition.

