## Appendix S1 for "Using spatial prioritization to navigate current and future tradeoffs when controlling invaders"

### Using spatial prioritization to navigate current and future tradeoffs when controlling invaders

Brendan K. Hobart, H. Anu Kramer, Julianna M. A. Jenkins, Damon B. Lesmeister, Raymond J. Davis, M. Zachariah Peery

Contents:

Table S1 (p. 2–3)

Table S2 (p. 4)

Table S3 (p. 5)

Figure S1 (p. 6)

Figure S2 (p. 7)

Table S1. Species whose geographic ranges were included as spatial inputs in our prioritization schemes. These taxa are all state or federally listed and putatively affected by barred owls through competition or predation. This list is an abbreviated version of Table 3-29 in the UWFWS Barred Owl Management Strategy Environmental Impact Statement. Some species in the original table were excluded from our analyses because they (1) did not occur within our study area and/or (2) did not have geographic range information available. Taxa included in Listing statuses are denoted as: E = endangered, T = threatened, C = candidate, PT = proposed threatened, PE = proposed endangered, S = species of concern (or similar).

| Common name | Scientific name | Fed status | WA status | OR status | CA status | Barred owl prey | Barred owl competitor |
|---|---|---|---|---|---|---|---|
| Great grey owl | *Strix nebulosa* | - | - | S | E | - | YES |
| Little willow flycatcher [1] | *Empidonax traillii brewsteri* | - | - | - | E | YES | - |
| Marbled murrelet | *Brachyramphus marmoratus* | T | E | E | E | YES | - |
| Oregon vesper sparrow [1] | *Pooecetes gramineus affinsis* | - | E | S | S | YES | - |
| California tiger salamander | *Ambystoma californiense* | E/T | - | - | T | YES | - |
| California red-legged frog | *Rana draytonii* | T | - | - | S | YES | - |
| Foothill yellow-legged frog | *Rana boylii* | PE/PT | S | S | E/T | YES | - |
| Oregon spotted frog | *Rana pretiosa* | T | S | S | S | YES | - |
| Scott Bar salamander | *Plethodon asupak* | - | - | - | T | YES | - |

| | | | | | | | |
|---|---|---|---|---|---|---|---|
| Shasta salamander | *Hydromantes shastae, H. samweli, H. wintu* | - | - | - | T | YES | - |
| Siskiyou mountains salamander | *Plethodon stormi* | - | - | S | T | YES | - |
| Western pond turtle | *Actinemys marmorata* | - | E | S | S | YES | - |
| Canada lynx | *Lynx canadensis* | T | E | - | - | - | YES |
| Cascade red fox [1] | *Vulpes vulpes cascadensis* | - | E | - | - | - | YES |
| Fisher | *Pekania pennanti* | E (partial) | E | S | T/S | - | YES |
| Mazama pocket gopher | *Thomomys mazama* | T | T | - | - | YES | - |
| Pacific (Humboldt) marten [2] | *Martes caurina humboldtensis* | T (partial) | - | S | E | - | YES |
| Point Arena mountain beaver | *Aplodontia rufa nigra* | E | - | - | S | YES | - |
| Red tree vole | *Arborimus longicaudus* | C | - | S | - | YES | YES |
| Sierra Nevada red fox | *Vulpes vulpes necator* | E | - | S | T | - | YES |
| Western grey squirrel | *Sciurus griseus* | - | T | S | - | YES | - |

[1] These taxa did not have subspecies-level range data available through the USGS GAP, and thus base species range data were used in our analyses
[2] At the time USGS GAP was compiled, the Pacific marten was conspecific with the American marten (*Martes americana*), and thus we used geographic range data for the former

Table S2. Weights used to design prioritization scenarios. Note that federally reserved forest and fire refugia were included as penalties with negative weight values to functionally incentivize their distribution in solutions.

| Threat | Spotted owl weight ($W_{NSO}$) | At-risk species weights* | Fire penalty parameter | Federally reserved forest penalty parameter | Fire refugia penalty parameter | Distance-to-road penalty parameter | Distance-to-trail penalty parameter |
|---|---|---|---|---|---|---|---|
| Current | 1 | 1 | 1 | -1 | -1 | 1 | 1 |
| Current | 5 | 1 | 1 | -1 | -1 | 1 | 1 |
| Current | 10 | 1 | 1 | -1 | -1 | 1 | 1 |
| Current | 15 | 1 | 1 | -1 | -1 | 1 | 1 |
| Future | 1 | 1 | 1 | -1 | -1 | 1 | 1 |
| Future | 5 | 1 | 1 | -1 | -1 | 1 | 1 |
| Future | 10 | 1 | 1 | -1 | -1 | 1 | 1 |
| Future | 15 | 1 | 1 | -1 | -1 | 1 | 1 |

*each at-risk species geographic range was included individually with a weight of 1

Table S3. Distribution of high-priority hexagons (i.e., those with relative importance scores >0.9) among regions, for each scenario.

| Threat | $W_{NSO}$ | region | % high priority cells |
|---|---|---|---|
| future | 1 | Klamath | 8.67 |
| | | Oregon & California Cascades | 78.11 |
| | | Oregon Coast | 13.22 |
| future | 5 | Klamath | 34.56 |
| | | Oregon & California Cascades | 64.56 |
| | | Oregon Coast | 0.89 |
| future | 10 | Klamath | 60.67 |
| | | California Coast | 23.33 |
| | | Oregon & California Cascades | 16.00 |
| future | 15 | Klamath | 51.67 |
| | | California Coast | 47.44 |
| | | Oregon & California Cascades | 0.89 |
| current | 1 | Klamath | 9.11 |
| | | Oregon & California Cascades | 73.78 |
| | | Oregon Coast | 17.11 |
| current | 5 | Klamath | 33.56 |
| | | Oregon & California Cascades | 62.22 |
| | | Oregon Coast | 4.22 |
| current | 10 | Klamath | 63.67 |
| | | California Coast | 3.22 |
| | | Oregon & California Cascades | 31.89 |
| | | Oregon Coast | 1.22 |
| current | 15 | Klamath | 79.11 |
| | | California Coast | 5.67 |
| | | Oregon & California Cascades | 14.67 |
| | | Oregon Coast | 0.56 |

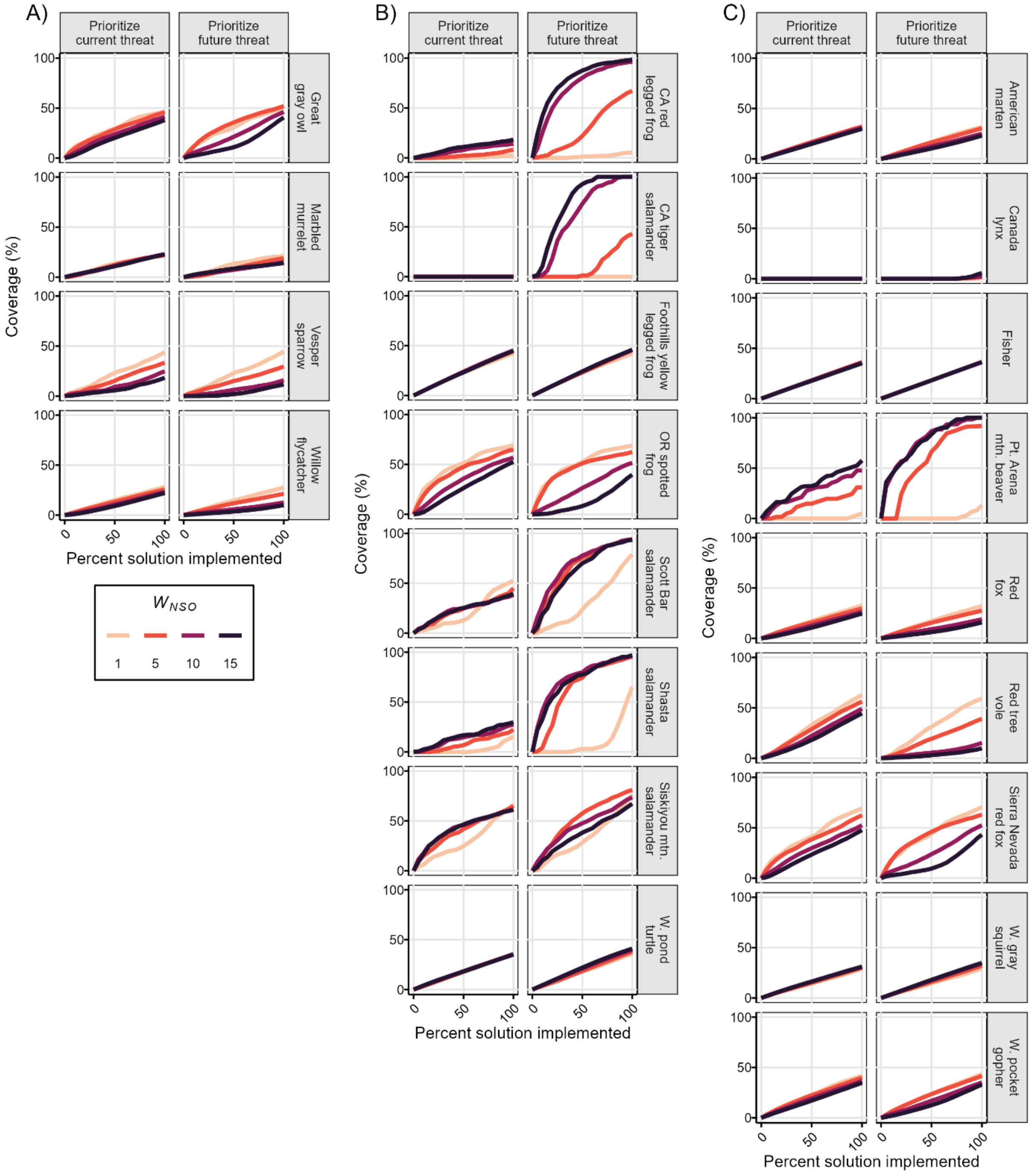
A)
B)
C)
Prioritize current threat
Prioritize future threat
Coverage (%)
Percent solution implemented
Great gray owl
Marbled murrelet
Vesper sparrow
Willow flycatcher
CA red legged frog
CA tiger salamander
Foothills yellow legged frog
OR spotted frog
Scott Bar salamander
Shasta salamander
Siskiyou mtn. salamander
W. pond turtle
American marten
Canada lynx
Fisher
Pt. Arena mtn. beaver
Red fox
Red tree vole
Sierra Nevada red fox
W. gray squirrel
W. pocket gopher
$W_{NSO}$
1
5
10
15

Figure S1. Individual species-level coverage plots. A curve that rises more steeply and to a greater maximum is indicative of a more efficient solution—greater coverage is afforded with relatively less effort (that is, less of the solution being implemented).

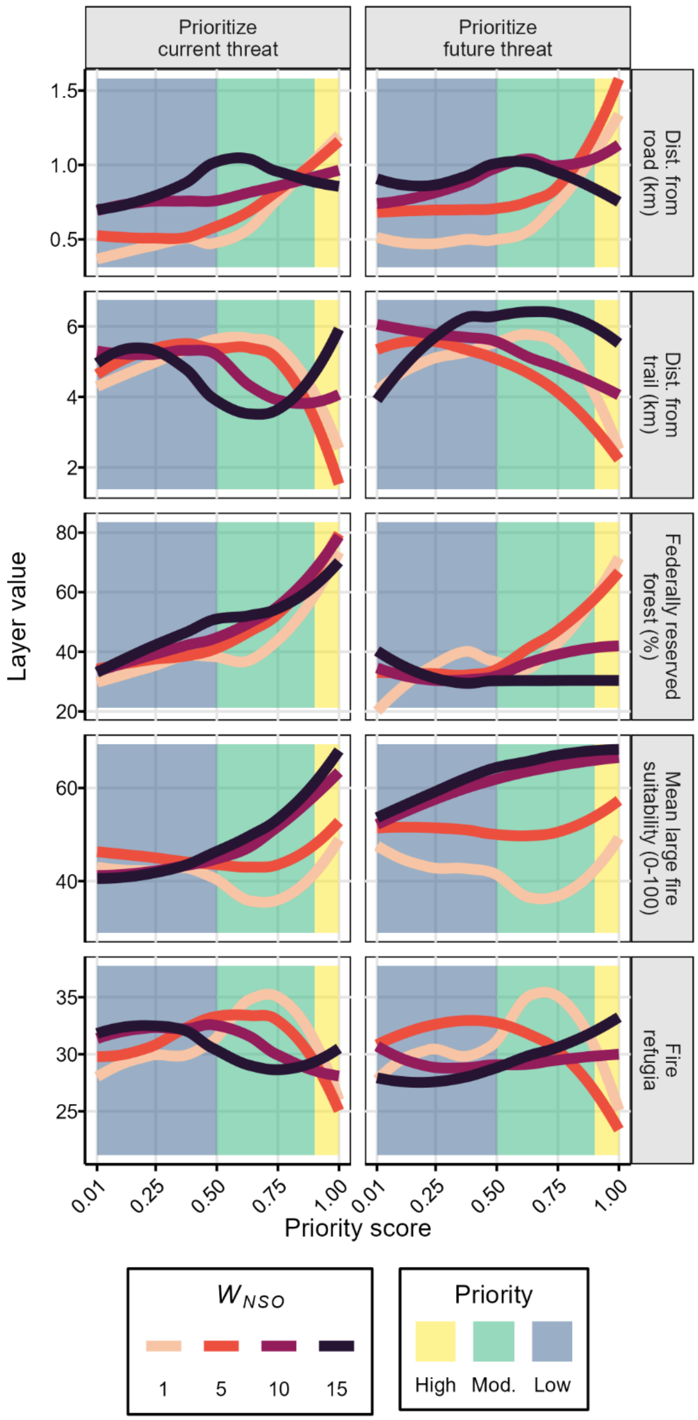


Figure S2. Smoothed distributions of input penalty layers across areas of low-to-high priority. We calculated mean layer values across all hexes within equal-interval priority score bins, then used basic general additive models to plot smoothed lines to mean values across priority scores.
